# Interleukin Enhancer Binding Factor 2 (Ilf2) and Kidney Epithelial Stress Resilience

**DOI:** 10.1101/2025.08.12.667599

**Authors:** Shuang Cao, Karen I. López-Cayuqueo, Janna Leiz, Felix J. Boivin, Lajos Markó, Christian Hinze, Jessica Schmitz, Jan Hinrich Bräsen, Kai M. Schmidt-Ott

## Abstract

**Background:** Epithelial cells in the renal medulla are continuously exposed to hyperosmolality, hypoxia, and oxidative stress, yet they display remarkable resilience. The transcriptional programs that endow this intrinsic stress tolerance remain incompletely defined.

**Methods:** We integrated single-nucleus RNA sequencing of mouse kidneys, computational transcription factor (TF) prioritization, and single-cell CRISPR interference screening (Perturb-seq) in inner medullary collecting duct (IMCD3) cells to systematically identify TFs mediating epithelial stress adaptation. The role of Ilf2 (Interleukin Enhancer Binding Factor 2) was further evaluated in IMCD3 cells by bulk RNA-seq, splicing analysis, functional assays under hyperosmotic stress, as well as in a mouse model of kidney ischemia–reperfusion injury (IRI) and kidney tissues from patients with early and advanced chronic kidney disease (CKD).

**Results:** Computational prediction and Perturb-seq identified known and novel TFs, that regulate gene expression programs in kidney medullary tubules. Among them, Ilf2 (also called NF45) emerged as a previously unrecognized regulator: Ilf2 knockdown in IMCD3 cells disrupted transcriptional and splicing programs linked to cell proliferation, cytoskeletal organization, and stress adaptation. Ilf2-deficient cells exhibited reduced proliferation, impaired nuclear integrity, and increased sensitivity to hyperosmotic stress. In mouse kidneys, Ilf2 expression increased during tubular repair after IRI, accompanied by induction of Ilf2-dependent transcripts and splicing events. Human kidneys with advanced CKD displayed elevated expression and cytoplasmic translocation of ILF2, suggesting a conserved stress-adaptive response.

**Conclusions:** ILF2 orchestrates both transcriptional and post-transcriptional regulatory mechanisms to sustain kidney epithelial stress resilience. Our findings highlight ILF2 as a potential tubular stress biomarker and therapeutic target for enhancing renoprotection.

**Key Points:**

1. Ilf2 has significant and profound impact on gene expression programs in cultured kidney medullary epithelial cells (IMCD3).
2. Ilf2 confers cellular resilience, nuclear integrity, RNA splicing, proliferative capacity, and osmotic resistance in IMCD3 cells.
3. Injured kidneys display increased levels of Ilf2 and up-regulation of genes and splicing events related to Ilf2 function.

## Introduction

The mammalian kidney maintains fluid and electrolyte balance through a complex cortico–medullary architecture. Urine concentration depends on countercurrent multiplication and urea recycling, which establish a hyperosmolar interstitium in the medulla to facilitate water reabsorption^1^. As a result, medullary collecting duct cells experience persistent osmotic, hypoxic, and oxidative stress^2,3^. Remarkably, these epithelial cells not only withstand such harsh conditions but also mount adaptive responses, a property that becomes particularly critical in injured kidneys, where collecting duct cells display greater injury resistance than proximal tubules^4,5^.

Transcription factors (TFs) orchestrate stress-response programs that preserve epithelial function^6–8^. In collecting duct principal cells, TFs such as Grainyhead-like 2 (GRHL2) and NFAT5 (TonEBP) safeguard epithelial integrity and osmoprotection, for example by maintaining tight junctions and inducing osmoprotective genes including Aqp2^9–11^. While additional collecting duct pathways, such as Yap and Tfcp2l1, have been implicated in collecting duct homeostasis and disease^12,13^, the broader transcriptional networks underlying medullary epithelial resilience remain poorly understood.

Systematic functional dissection of candidate TFs identified from omics datasets has been constrained by the low throughput and labor-intensive nature of traditional knockout approaches. Recent advances in pooled CRISPR interference (CRISPRi) combined with single-cell RNA sequencing (scRNA-seq) now enable parallel interrogation of multiple regulators at cellular resolution, providing a powerful means to dissect molecular mechanisms^14,15^.

To systematically identify transcriptional regulators of medullary tubular stress resilience, we combined *in vivo* and *in vitro* functional genomics. We first inferred cell-type–specific regulatory networks from mouse kidney single-nucleus RNA-seq data and prioritized TFs with high activity in medullary collecting duct cells. Using CRISPRi Perturb-seq in IMCD3 cells, we then conducted parallel functional screening at single-cell resolution. This integrative strategy uncovered Interleukin Enhancer Binding Factor 2 (ILF2/NF45) as a previously unrecognized regulator of epithelial stress adaptation. Although ILF2 has been characterized in cancer and immune regulation, its function in kidney tubules has remained unknown. Here, we show that ILF2 safeguards epithelial proliferation, nuclear integrity, and osmotic-stress tolerance by coordinating transcriptional and alternative-splicing programs. These findings identify ILF2 as a central integrator of transcriptional and post-transcriptional control that reinforces kidney epithelial resilience under stress.

## Methods

### Animals

All animal experiments were approved by and performed in compliance with local authorities (LAGeSo Berlin, Germany). Mice were housed under standard conditions in the Max-Delbrück-Centrum für Molekulare Medizin (MDC) animal facility according to institutional guidelines. Experiments were performed in 8- to 12-week-old males.

### Mouse Kidney snRNA-seq Sample Preparation

Following euthanasia, from each of the two wild-type (C57BL/6) mice, a single kidney (left or right, selected randomly) was harvested and immediately placed in ice-cold PBS. A 1–2 mm coronal section containing cortex, outer medulla, and inner medulla was isolated based on a previously described protocol (Leiz et al., J Vis Exp, 2021)^16^. Tissue samples were incubated in RNAlater at 4°C for 24 hours, then stored at – 80°C until further processing.

For nuclei isolation, frozen samples were thawed, minced in nuclear lysis buffer (NLB1) supplemented with 1 U/μl RNase inhibitor and 10 mM ribonucleoside vanadyl complex (RVC), and homogenized using a Dounce homogenizer. The suspension was filtered sequentially through 100 μm, 40 μm, and 20 μm strainers, with centrifugation and sucrose cushion steps to enrich nuclei. The resulting nuclei pellet was washed and resuspended in PBS containing 0.04% BSA and 1 U/μl RNase inhibitor, stained with DAPI. Single nuclei were sorted by FACS (BD FACS Aria II) to remove debris and multiplets. Approximately 10,000 nuclei per sample were loaded into the 10X Genomics Chromium platform (v3.1) for snRNA-seq.

### snRNA-seq Data Processing and Analysis

Raw sequencing data were processed using Cell Ranger (v3.0.2, 10X Genomics) to align reads and generate gene-cell matrices. Filtered feature-barcode matrices were imported into Seurat (v4.4.3)^17^ for downstream analysis. Nuclei with fewer than 600 or more than 8000 detected genes, or with >2.5% mitochondrial reads, were excluded from further analysis.

Dimensionality reduction and clustering were performed using the RunPCA, FindNeighbors (dims = 1:30), FindClusters (resolution = 0.5), and RunUMAP functions in Seurat. Marker genes were identified using the FindAllMarkers function with default parameters. A total of 13 transcriptionally distinct clusters were identified and annotated based on canonical marker gene expression.

### Candidate Transcription Factors Selection

Single-nucleus RNA-sequencing data from the kidneys of two wild-type (C57BL/6) mice were used to infer transcriptional regulatory networks across all cell types. Regulatory network inference was performed using the Single-Cell rEgulatory Network Inference and Clustering (SCENIC) pipeline, implemented via the pySCENIC framework (v0.11.2)^19^. The analysis comprised three steps: (1) gene regulatory network (GRN) inference based on co-expression, (2) cis-regulatory motif enrichment and pruning (ctx), and (3) regulon activity scoring (AUCell). The mm10 reference genome was used throughout the analysis. Motif enrichment was performed using the cisTarget v9 motif database. Specifically, the following resources were employed: the gene-motif ranking file mm10 refseq-r80 10kb_up_and_down_tss.mc9nr.genes_vs_motifs.rankings.feather, the motif annotation file motifs-v9-nr.mgi-m0.001-o0.0.tbl, and a curated list of mouse transcription factors (allTFs_mm.txt). These resources were obtained from the cisTarget and SCENIC resource portal, maintained by the Aerts Lab. Default parameters were applied unless otherwise specified.

SCENIC computes regulon specificity scores (RSS) and regulon activity scores (RAS) of each cell type cluster to quantify the regulatory potential of transcription factors (TFs) across single-cell populations **(Supplementary Table S2)**. To prioritize candidate TFs for functional validation, RSS and RAS values from the medullary collecting duct principal cells cluster (OMCD and IMCD) were integrated with transcriptomic data from wild-type IMCD3 cells (n = 3, bulk RNA-seq, **Supplementary Table S3**) and normalized gene expression values derived from snRNA-seq. Candidate TFs were selected based on the following criteria: RAS > 0.5, RSS > 0.05, IMCD3 transcript-per-million (TPM) > 10, and average log-normalized expression in CD-PC cells > 0.5. The full list of candidate TFs and corresponding selection metrics is summarized in **Supplementary Table S1**.

### Cell Culture and sgRNA Lentivirus Library Transduction

Mouse inner medullary collecting duct (IMCD3) cells (ATCC CRL-2123, Manassas, USA) were cultured in DMEM + GlutaMAX (Gibco, 61965059) supplemented with 10% fetal bovine serum (FBS; Gibco, A5256701) and penicillin-streptomycin (100 U/mL and 100 µg/mL, respectively; Gibco, 15140122). Cells were maintained at 37°C with 5% CO_₂_ and passaged at a 1:4 ratio upon reaching 90% confluency using 0.25% trypsin (Gibco, 15090046) in PBS. To generate dCas9-KRAB-expressing IMCD3 cells, cells at 50% confluency were transduced with lentiviral particles encoding KRAB-dCas9 (10X CRISPRi Feature Barcode Optimization Kit; Sigma-Aldrich) at a multiplicity of infection (MOI) of 1. After 24 hours, the transduction medium was replaced with a fresh culture medium, and cells were allowed to recover for an additional 24 hours. Blasticidin (1 µg/mL final concentration) was then added for selection, which was maintained for 7 days. The stable expression of dCas9-KRAB was confirmed by western blot (**Supplementary Figure S1**).

For single-cell CRISPR (sc-CRISPR) screening, we first performed a pilot experiment to confirm the efficiency of the gRNA capture vector **(Supplementary Figure S2)**. A lentiviral library of guide RNAs (gRNAs; designed and packaged by Sigma-Aldrich) was then transduced into dCas9-KRAB IMCD3 cells at 70% confluency with a multiplicity of infection (MOI) of 0.5. After 24 hours, cells were recovered in fresh medium for an additional 24 hours and subsequently selected with 1 µg/mL puromycin for 7 days to remove un-transduced cells.

### Western Blotting

Total protein was extracted from cells using RIPA lysis buffer (Sigma, R0278-50ML) supplemented with one tablet of protease inhibitor cocktail (Roche, 11836170001) and PhosSTOP phosphatase inhibitor (Roche, 04906845001). Lysates were centrifuged at 13,000 × g for 30 minutes at 4 °C, and the supernatant was collected. Protein concentrations were determined using the BCA assay (Thermo Scientific, 23225).

Equal amounts of protein were separated by SDS-PAGE on 4%–12% Bis-Tris gels (Invitrogen, NP0322BOX) and transferred onto nitrocellulose membranes (Thermo Scientific, 88018). Membranes were blocked and incubated with primary antibodies at 1:1000 dilution unless otherwise specified. The following primary antibodies were used: anti-Cas9 (Sigma, MAC133), anti-Ilf2 (Abcam, ab113205), and anti-Vinculin (Proteintech, 66305-1-Ig). Band intensities were quantified using ImageJ2 software.

### Single-cell CRISPR Screen Libraries Construction and Sequencing

On the day of single-cell capture, successfully transduced dCas9-KRAB IMCD3 cells were harvested and dissociated using 0.25% trypsin to generate a single-cell suspension. The suspension was filtered through a 30 µm cell strainer (MACS SmartStrainers, 130-098-458) to ensure single-cell separation. Viable cells were counted using a Countess II Automated Cell Counter (Thermo Fisher) with Trypan Blue exclusion. All procedures were performed following the 10x Genomics Cell Preparation Demonstrated Protocol (CG00054, Rev B).

Single-cell encapsulation was performed on the 10x Chromium Controller, targeting 10,000 cells per channel. Gene expression and CRISPR guide RNA libraries were constructed using the Chromium Single Cell 3D Reagent Kits v3 (10x Genomics) with Feature Barcode technology for CRISPR screening (CG000184, Rev C), following the manufacturer’s instructions.

The resulting libraries were sequenced on an Illumina NovaSeq 6000 platform using paired-end mode. Raw sequencing data were processed using the Cell Ranger software (v7.1.0) with the CRISPR Guide Capture Analysis pipeline. Reads were aligned to the mouse mm10 genome (GENCODE vM23, Ensembl 98), and digital expression matrices were generated for downstream analysis.

### CRISPR Screening Data Analysis

The filtered_feature_bc_matrix output from Cell Ranger was imported into Seurat (version 4.3.0) for downstream analysis. Cell quality was assessed based on total UMI counts and proportion of mitochondrial gene reads, quality control metrics are provided in **Supplementary Figure S3**. Cells were assigned to the perturbation group based on their captured guide RNA (gRNA) sequences derived from Feature Barcode data. Perturbation efficiency and transcriptomic responses were evaluated by grouping cells according to their associated gRNAs. Differential gene expression analysis between defined cell populations was performed using the DESeq2 package^20^.

### Functional Enrichment Analysis of Differentially Expressed Genes

Gene Ontology (GO) and KEGG-based enrichment analyses were performed using the ClusterProfiler package (v4.14.6) in R. Differentially expressed genes (DEGs) were identified using DESeq2 with an adjusted p-value < 0.05. GO term enrichment including Biological Process (BP), Molecular Function (MF), and Cellular Component (CC) was assessed using the enrichGO() function.

For KEGG pathway analysis, a curated subset of pathways related to metabolism, signal transduction, and cellular processes was selected from the KEGG PATHWAY database for *Mus musculus*^21^. Gene sets corresponding to these pathways were compiled and used as custom input to the enricher() function. The gene universe was defined as all expressed genes in the dataset. Multiple testing correction was applied using the Benjamini-Hochberg (BH) method, and terms with adjusted p-value < 0.05 were considered statistically significant. Results were visualized using ggplot2.

### Gene Set Enrichment Analysis

Gene set enrichment analysis (GSEA) was performed using the GSEA desktop software (v4.3.3) to assess pathway-level enrichment. Normalized gene expression data were obtained via DESeq2 from single-cell RNA-seq. Hallmark gene sets from the Molecular Signatures Database (MSigDB v2024.1, mouse version) were used as the reference collection (mh.all.v2024.1.Mm.symbols.gmt). GSEA was run in gene set permutation mode using the t-test statistic as the ranking metric, with 1,000 permutations and gene set size thresholds set to a minimum of 10 and maximum of 500 genes. Enrichment scores were calculated using the weighted scoring scheme, and pathways with an adjusted p-value < 0.05 were considered significantly enriched.

### Generation and Validation of the Stable Ilf2 knockdown IMCD3 Cell Line

Three guide RNAs (gRNAs) targeting Ilf2, previously used in the screening experiment, were selected to generate a stable Ilf2-knockdown IMCD3 cell line. The selected gRNAs were cloned into a lentiviral expression vector (pLV[gRNA]-EGFP:T2A:Puro-U6; VectorBuilder) and packaged into lentiviral particles. dCas9-KRAB IMCD3 cells were transduced with the resulting lentiviruses to induce CRISPR interference-mediated Ilf2 knockdown. Knockdown efficiency was validated at both the transcript and protein levels by quantitative PCR (qPCR) and western blot analysis, respectively. The primers used for Ilf2 detection were as follows:

Forward: 5’-CCTGGGGAACAAAGTCGTGG-3’

Reverse: 5’-TGAGAATTTTCACCGTAGCATCA-3’

### Quantitative PCR

Total RNA was extracted using the RNeasy Mini Kit (QIAGEN) according to the manufacturer’s instructions. RNA concentrations were quantified using a NanoPhotometer (N60; Implen). Complementary DNA (cDNA) was synthesized from 1 µg of total RNA using the RevertAid First Strand cDNA Synthesis Kit (Thermo Fisher Scientific). Quantitative real-time PCR (qRT-PCR) was performed on a LightCycler 96 System (Roche). Gene expression levels were normalized to GAPDH as an internal control, and relative expression was calculated using the 2^–ΔΔCt method.

The primers used for GAPDH detection were as follows:

Forward: 5’-AGCTTGTCATCAACGGGAAG-3’

Reverse: 5’-TTTGATGTTAGTGGGGTCTCG-3’

### Renal Ischemia-Reperfusion Model

Five male mice, aged 8 to 12 weeks, were anesthetized with isoflurane (2.3%) in air (350 ml/min) and kept on a warm pad at 37°C (Physitemp Instruments, Clifton, NJ, USA) to induce warm ischemia. Body temperature was monitored via a rectal probe. The left renal pedicle was exposed by an abdominal incision and clipped for 25 minutes (min) using a non-traumatic aneurysm clip (Aesculap, Berlin, Germany). Successful clamping of the left renal pedicle and the reperfusion (i.e., color change after removal of the clamp) was confirmed visually. After surgery, each mouse was subcutaneously injected with sterile physiologic saline solution (0.9% sodium chloride, 10 ul/g) and analgesic buprenorphine (0.1 mg/kg). Between the first pre-and third postoperative days, mice were provided with raspberry-flavoured metamizole (800 mg/kg/24 h) via drinking water for pain prophylaxis. The animals were sacrificed after 7 days. Both the contralateral and ischemic kidneys were collected for further analysis. Animals were monitored daily for general condition and signs of distress, including reduced mobility, weight loss, or abnormal posture. No animals reached humane endpoints before the planned sacrifice on day 7.

### Bulk RNA sequencing and splicing analysis

Bulk RNA sequencing was performed as an orthogonal validation to support the transcriptional effects observed in the single-cell CRISPR perturbation screen. Specifically, the aim was to assess whether Ilf2 knockdown in bulk IMCD3 cells could recapitulate gene expression changes identified in the single-cell context.

Total RNA was extracted from IMCD3 cells as described in the qPCR section. Library construction and paired-end sequencing (150 bp) were carried out by Novogene (Munich, Germany) using a poly(A)-enrichment protocol. Raw sequencing reads were quality-checked using FastQC (v0.11.8), no trimming was performed, as the sequencing quality was deemed sufficient. Clean reads were aligned to the mouse reference genome (*Mus musculus*, GRCm39, Ensembl release 110) using STAR (v2.7.11b), and gene-level quantification was performed with featureCounts (v2.0.6).

Differential expression analysis between control and Ilf2-knockdown cells (n = 6 biological replicates) was performed using the ComBat-seq^22^ approach to correct for batch effects. Alternative splicing analysis was performed using the R package SpliceWiz (v1.8.0), based on STAR-aligned BAM files. Functional enrichment analysis of differentially spliced genes (FDR < 0.05 and |ΔPSI| > 0.05) was conducted using clusterProfiler (v4.14.6) and visualized with ggplot2.

Alternative splicing analysis was performed using the R package SpliceWiz^23^ (v1.8.0), based on STAR-aligned BAM files. Functional enrichment analysis of differentially spliced genes (FDR < 0.05 and |ΔPSI| > 0.05) was conducted using clusterProfiler (v4.14.6) and visualized with ggplot2.

### In situ hybridization

Kidney tissues were fixed in 4% paraformaldehyde (PFA) in phosphate-buffered saline (PBS), embedded in paraffin, and sectioned at 5 μm thickness. Sections were mounted on Superfrost Plus slides (Thermo Scientific) and air-dried overnight.

Fluorescent in situ hybridization was performed using the RNAscope Multiplex Fluorescent Reagent Kit v2 (ACD, #323100) according to the manufacturer’s instructions. Formalin-fixed, paraffin-embedded (FFPE) kidney sections were hybridized with probes targeting Ilf2 (custom-designed, ACD) and Aqp4 (ACD, #417161-C2). Sections were counterstained with DAPI and mounted using Dako Fluorescence Mounting Medium (Dako, #S3023). Images were acquired using a Zeiss LSM 980 confocal microscope equipped with Airyscan 2.

### Immunofluorescence staining

Paraffin-embedded mouse and human kidney tissue sections were cut and deparaffinized through a graded ethanol series, followed by antigen retrieval in a Tris-EDTA buffer (pH 9.0). Immunohistochemistry was performed using an anti-Ilf2 antibody (Abcam, #ab113205; 1:500). A corresponding secondary antibody was applied at 1:1000 dilution. For co-staining, an anti-Aqp2 antibody (Santa Cruz, #sc-9882; 1:200) and its respective secondary antibody (1:1000) were used. In mouse kidney samples, Lotus Tetragonolobus Lectin (LTL, 1:1000) was used to label the proximal tubule brush border, whereas in human kidney sections, an anti-Lrp2 antibody (BiCell, #31012; 1:200) with its corresponding secondary antibody (1:1000) was applied for proximal tubule staining.

For immunofluorescence staining of cultured cells, samples were fixed with 4% paraformaldehyde (PFA) for 15 minutes at room temperature, followed by PBS washes. Cells were permeabilized with 0.1% Triton™ X-100 in 0.5% BSA/PBS for 15 minutes. A mouse anti-Ki67 antibody (Abcam, #ab15580; 1:1000 dilution) was used for proliferation staining. Actin filaments were labelled with Alexa Fluor™ 532 Phalloidin (Invitrogen) according to the manufacturer’s instructions. Nuclei were counterstained with DAPI (2 μg/mL). Fluorescence images were acquired using a Zeiss LSM 980 confocal microscope.

### Immunofluorescence quantification

Automated cell detection and Ki67-positive cell quantification were performed using QuPath (version 0.4.2) software. Regions of interest (ROIs) were manually defined if needed, and cell segmentation was based on nuclear DAPI signal. Cells were classified into Ki67-positive or Ki67-negative populations using custom-defined intensity thresholds or pre-trained classifiers. Batch processing was applied to all images to ensure consistency and reduce operator bias. Quantitative outputs including the number of Ki67-positive cells and total DAPI-positive cells were exported for further statistical analysis. Manual quantification of multinucleated cells was performed using ImageJ2. F-actin staining was used to delineate individual cell boundaries, and cells containing more than one DAPI-positive nucleus were identified as multinucleated using the Cell Counter plugin. Quantification of ILF2 expression in the mouse and human slides were carried out using QuPath. Nuclei were segmented using DAPI staining, and the cytoplasmic area of each cell was defined by expanding the nuclear boundary by 5 µm. Mean ILF2 fluorescence intensity was measured within the nuclear and cytoplasmic regions, respectively.

### Histological staining

Paraffin-embedded kidney sections (4 µm) were deparaffinized and rehydrated through a graded ethanol series. Periodic acid–Schiff (PAS) staining was performed following standard protocols to evaluate tubular and glomerular morphology and injury.

For fibrosis assessment, adjacent sections were stained with Sirius Red solution according to standard procedures. Histological images were acquired under bright-field microscopy. Collagen-positive areas were quantified by our pathology department.

### Hyperosmotic treatment and XTT assay in IMCD3 cells

IMCD3 cells were cultured in standard DMEM medium (baseline osmolality ∼300 mOsm/kg) supplemented with 10% fetal bovine serum (FBS). Cells were seeded into 6-well or 96-well plates one day prior to hyperosmotic treatment, depending on the intended downstream assay (bright-field imaging or XTT assay). Hyperosmolarity was induced by adding 150 mM NaCl to the culture medium, raising the final osmolality to approximately 600 mOsm/kg. Both control and Ilf2-knockdown cells were treated with 600 mOsm/kg medium for 12 or 24 hours. Bright-field images were captured at 12 h and 24 h.

Cell death and survival were assessed using ImageJ2 software. Images were processed using a standardized analysis pipeline, which included local contrast enhancement (CLAHE: block size = 127, histogram bins = 256, maximum slope = 3), background subtraction (rolling ball radius = 50 pixels), automatic thresholding, and particle analysis (size filter: 200–5,000,000 pixels²). The analysis was performed in batch mode to ensure consistency across samples. At the 12-hour time point, cell viability was assessed using the XTT assay (Thermo Fisher Scientific, X12223) according to the manufacturer’s instructions. Absorbance was measured at 450 nm and 660 nm (reference/background) to evaluate metabolic activity.

### Human sample collection

Archived formalin-fixed, paraffin-embedded (FFPE) kidney specimens from patients with mild or advanced chronic kidney disease (CKD) were obtained from the Department of Pathology, Hannover Medical School (MHH). All samples were anonymized prior to use. The study was approved by the Ethics Committee of MHH (approval no. 10666_BO_K_2022 and 10863_BO_K_2023) and conducted in accordance with the Declaration of Helsinki and institutional guidelines for research involving human tissue.

### Statistics

Statistical analyses were performed using GraphPad Prism 10 or R packages. For comparisons between two groups, either Welch’s t-test (for normally distributed data with unequal variances) or the Mann–Whitney U test (for non-normally distributed data) was applied, with p-values adjusted for multiple testing using the Benjamini–Hochberg correction when applicable. For comparisons among multiple groups, one-way ANOVA followed by Tukey’s multiple comparison test was used. Unless otherwise indicated, data are presented as mean ± standard deviation (SD). Sample size (n) refers to the number of biological replicates and is specified in the corresponding figure legends. A *P* value or adjusted *P* value of < 0.05 was considered statistically significant.

## Results

### Transcription factor prioritization and CRISPRi perturbation screen in IMCD3 cells

To establish a suitable cellular model for dissecting transcriptional regulation in the epithelial cells of the renal medulla, we cultured IMCD3 cells (an immortalized cell line derived from mouse renal medullary epithelial cells) and obtained scRNA-seq data, which we then mapped to native mouse kidney cell snRNA-seq data. Cultured IMCD3 cells predominantly mapped to native kidney cell types representing medullary tubular cells of the kidney, including medullary collecting ducts (MCD) and thin limbs (TL) of Henle, supporting their use as an *in vitro* model that captures key transcriptional features of these cell types **(Figure 1a)**. We next sought to systematically identify candidate transcription factors (TFs) governing MCD cell identity and function. Using SCENIC-based regulon analysis^18^ of mouse kidney snRNA-seq data and filtering for TFs highly expressed in both native mouse kidney MCD cells and in cultured IMCD3 cells, we prioritized 25 high-confidence TFs for functional screening **(Figure 1b)**. To interrogate these TFs in a pooled manner, we implemented CRISPRi Perturb-seq^14^ in IMCD3 cells, enabling direct measurement of knockdown efficacy and transcriptomic consequences at single-cell resolution. The screen achieved broad coverage across gRNAs **(Figure 1c, Supplementary Figure S4)** and demonstrated robust suppression of target gene expression for the majority of TFs **(Figure 1d)**. Differential expression analysis showed that most TF perturbations induced modest transcriptomic changes **(Figure 1e)**. As a predefined positive control, Grhl2 knockdown reproduced the expected broad regulatory program, with >1,400 genes significantly altered and canonical downstream targets such as Cldn4, Epcam, and Rab25 robustly downregulated, consistent with the published literature^10^ **(Figure 1e–h, Supplementary Table S4)**. This recapitulation of known Grhl2-dependent effects demonstrates the robustness of our CRISPRi Perturb-seq platform. Notably, among the remaining candidates, Ilf2 perturbation produced the second largest transcriptomic shift, nominating Ilf2 as a previously unrecognized regulator of renal medullary epithelial cell programs and motivating further mechanistic investigation in subsequent analyses.

**Figure 1.**
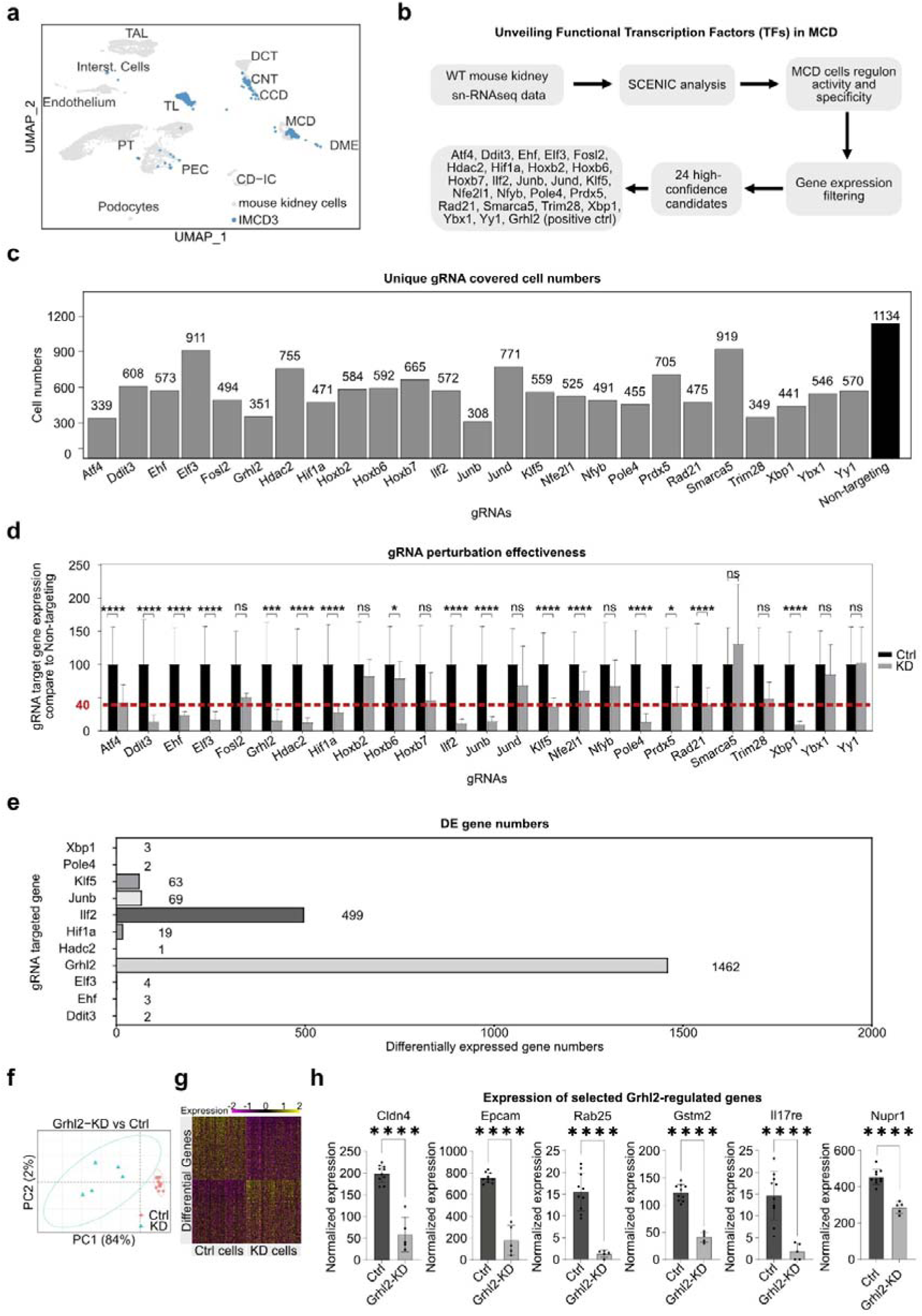
Identification of functional transcription factors in medullary collecting duct cells (MCD) by single-cell CRISPRi Perturb-seq. (**a**) Single cell transcriptomes of IMCD3 cells were projected onto a UMAP embedding of mouse kidney snRNA-seq reference data using MapQuery approach and reference label transfer. PT, proximal tubule; TAL, thick ascending limbs; TL, thin limbs; PEC, parietal epithelial cells; DCT, distal convoluted tubules; CNT, connecting tubules; CCD, cortical collecting ducts; MCD, medullary collecting ducts; DME, deep medullary epithelium; CD-IC, collecting duct intercalated cells. (**b**) Workflow for selecting the candidate TFs, regulon activity and specificity were inferred from wild-type mouse kidney snRNA-seq data using SCENIC (RcisTarget + AUCell). Candidate TFs with high activity in MCD and detectable expression (TPM > 10) in both MCD and IMCD3 cells were filtered, yielding 25 high-confidence TFs for functional screening by CRISPRi Perturb-seq. (**c**) Bar plots show the number of high-quality single cells uniquely assigned to each gRNA (after QC filtering for single gRNA identity, mitochondrial read percentage, and UMI counts). Each TF was targeted by 5 gRNAs; and the non-targeting targeted by 10 gRNAs. (**d**) Quantification of sgRNA-targeted TF mRNA expression (KD) relative to non-targeting controls (Ctrl). (**e**) For each targeted TF, DE genes were identified relative to NT cells (FDR < 0.05). (**f**) PCA revealed clear segregation of Grhl2_KD and control cells. (**g**) Normalized expression of significantly DE genes comparing Grhl2 knockdown with control cells. (**h**) Expression of selected Grhl2-regulated genes. Normalized expression levels (mean ± SD) of representative targets. Pseudo-bulk differential expression analysis was performed using DESeq2. Gene-wise significance was determined using a Wald test on the negative binomial model, and p-values were adjusted for multiple comparisons using the Benjamini-Hochberg method within DESeq2. * adjusted *P* < 0.05; ** adjusted *P* < 0.01; *** adjusted *P* < 0.001, **** adjusted *P* < 0.0001.

### CRISPRi Perturb-seq implicates Ilf2 in transcriptional control of epithelial stress-response programs

To profile Ilf2-dependent transcriptional programs, we assessed per-guide representation versus single-cell coverage as a quality control metric. Non-targeting guides displayed the expected strong positive correlation between sequencing read counts and cell numbers, indicating uniform sampling of the pooled library **(Figure 2a, right)**. By contrast, Ilf2-targeting guides deviated from this pattern (r = –0.63, p = 0.26; **Figure 2a, left**), suggesting reduced fitness or survival of Ilf2-deficient cells. PCA and gene-level heatmaps revealed clear separation of Ilf2-knockdown cells from controls, consistent with widespread transcriptomic remodeling **(Figure 2b–c)**.

**Figure 2.**
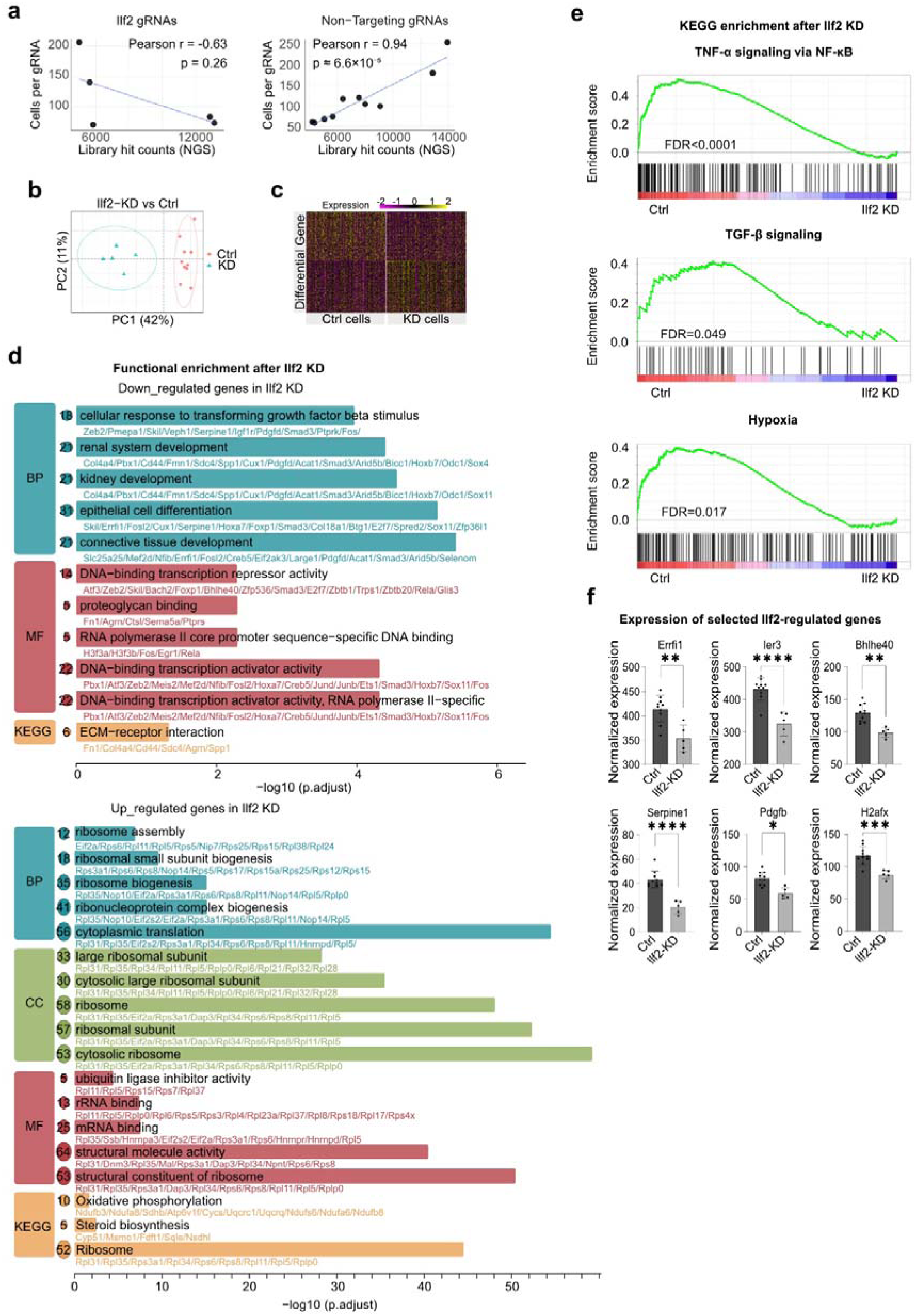
Transcriptional consequences of Ilf2 knockdown in IMCD3 cells revealed by Perturb-seq. **(a)** Scatter plots show correlation of gRNA abundance between the initial lentiviral library (NGS counts, x-axis) and captured gRNA in single cells (y-axis). For Ilf2-targeting gRNAs, library-to-cell correlation was moderate and negative (Pearson r = -0.63, p = 0.26), whereas non-targeting controls showed strong correlation (Pearson r = 0.94, p = 9.6 × 10⁻L). **(b)** PCA of single-cell transcriptomes demonstrates clear separation between Ilf2-KD and non-targeting control cells, with PC1 and PC2 explaining 42% and 11% of variance, respectively. **(c)** Heatmap of Ilf2-regulated genes. Z-scored expression of differentially expressed (DE) genes (FDR < 0.05). **(d-e)** Functional enrichment of Ilf2-dependent genes. GO (BP, CC, MF) and KEGG enrichment analysis of DE genes. **(f)** Expression of selected DE genes (mean ± SD) in Ilf2-KD versus control cells. Pseudo-bulk differential expression analysis was performed using DESeq2^20^. Gene-wise significance was determined using a Wald test on the negative binomial model, and p-values were adjusted for multiple comparisons using the Benjamini-Hochberg method within DESeq2. *adjusted *P* < 0.05; ** adjusted *P* < 0.01; *** adjusted *P* < 0.001, **** adjusted *P* < 0.0001.

Functional enrichment of differentially expressed genes showed that Ilf2 loss attenuates adaptive epithelial programs, including cellular response to TGF-β, extracellular-matrix organization, and transcriptional regulation while promoting ribosome biogenesis and translation-related pathways **(Figure 2d, Supplementary Table S5)**. GSEA further confirmed modulation of canonical stress-response pathways such as NF-κB, TGF-β, and hypoxia signaling **(Figure 2e)**. Representative stress-associated genes, including Errfi1, Ier3, Bhlhe40, Serpine1, Pdgfb, and H2afx, were all downregulated upon Ilf2 knockdown **(Figure 2f, Supplementary Table S4)**. Together, these findings indicate that Ilf2 maintains the transcriptional architecture that enables collecting duct epithelial cells to adapt to environmental stress.

### Bulk RNA-seq confirms Ilf2-dependent programs and reveals widespread alternative splicing

To orthogonally validate the screen, we generated an independent Ilf2 CRISPRi knockdown (KD) model in IMCD3 cells utilizing three different guide RNAs. ILF2 protein and mRNA were robustly reduced across guides **(Figure 3a–c)**. Bulk RNA-seq recovered a subset of the screen-derived DE genes **(Figure 3d and e)** and showed strong replicate concordance **(Figure 3f)**. Consistent with the pathway- and gene-level predictions from the screen **(Figure 2d-e)**, representative stress-reaction genes were down-regulated upon Ilf2 KD, including Serpine1, Pmepa1, Fosl2, Errfi1, Nasp, Sdc4, Sox4, Farp1, Ier3, H2afx, Larg1, Ext1, Cux1, Fn1 and Pdgfb **(Figure 3g, Supplementary Table S6)**.

**Figure 3.**
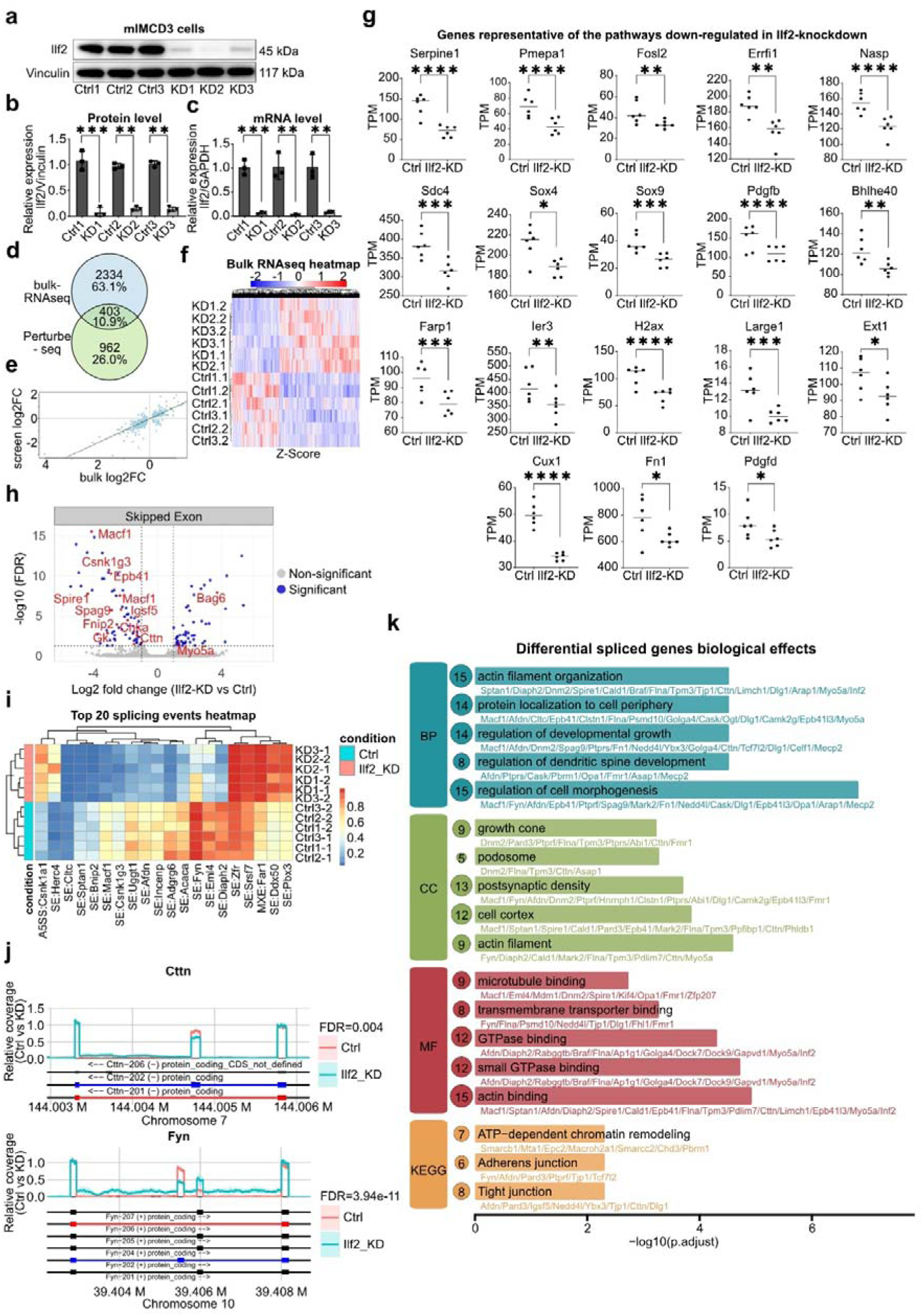
Bulk RNA-seq validation and alternative splicing changes upon Ilf2 knockdown. **(a)** Immunoblotting was conducted on cells from three independent knockdown models, using 3 different gRNA (KD1-3). Vinculin (Housekeeping protein) and Ilf2 were detected, verifying the efficacy of the knockdown. **(b)** The intensity of the protein bands was quantified using ImageJ. This quantification involved measuring the signal intensity of the bands to evaluate the relative protein levels in the knockdown versus control cells (Ctrl n=3). A t-test with Welch’s correction was performed on three independent experiments to determine statistical significance. **(c)** Quantitative PCR (qPCR) was performed on the established Ilf2 knockdown IMCD3 cell model using two different non-targeting gRNAs and two Ilf2 gRNAs. Quantification was calculated for both control cells and Ilf2 knockdown cells (n=3). A t-test with Welch’s correction was performed on three independent experiments to determine statistical significance. **(d)** Venn diagram shows overlap of DEGs (*P* < 0.05) detected by Perturb-seq and bulk RNA-seq. **(e)** Scatter plot illustrates correlation of log2 fold changes between the screening ScRNA-seq and bulk RNA-seq, the green line shows the actual correlation, and the dashed line shows the ideal fit. **(f)** Heatmap showing Z-score normalized expression of differentially expressed genes (DEGs) across biological replicates of Ilf2 knockdown (KD) and control (Ctrl) groups from 3 different gRNA mediated Ilf2 knockdowns and controls (n=6). Rows represent genes, while columns represent individual samples. Red indicates upregulated genes, and blue indicates downregulated genes, relative to the mean expression across samples. **(g)** Expression levels of selected genes were analyzed for validation purposes, comparing Ilf2 KD versus Ctrl (n = 6 per group, comprising 3 non-targeting and 3 Ilf2-targeting gRNAs). Expression values were normalized to transcripts per million (TPM) for visualization. Genes shown were selected based on their involvement in the pathways highlighted in figure 2. Differential expression analysis was performed on raw count data using DESeq2, with batch effects adjusted using ComBat-seq^22^. **(h)** Volcano plots showing differential splicing events (skipped exon) in Ilf2-knockdown (KD) versus control (Ctrl) cells. The x-axis indicates log_₂_ fold change in event inclusion; the y-axis indicates –log_₁₀_(FDR). Multiple testing correction was performed using the Benjamini–Hochberg correction to determine statistical significance. In the plot, genes with an adjusted P < 0.05 and a log_₂_ fold change > 1 were considered significant. Selected genes are labeled in red (compare to Fig. 5 below). **(i)** Heatmap of the top 20 most significantly altered splicing events, based on PSI (Percent Spliced-In) values. Rows indicate biological replicates, columns represent individual events; red and blue denote higher and lower inclusion levels, respectively. **(j)** Representative examples of alternative splicing changes in Cttn and Fyn. Coverage plots display normalized read densities across the Cttn (chr7:144.003–144.006 Mb) and Fyn (chr10:39404–39408 Mb) gene loci. Each curve represents the mean normalized coverage across replicates for each condition: control (red) and KD (blue). Shaded areas indicate the standard error. Transcript isoforms are shown below with exon–intron structures. **(k)** Enrichment Combined GO and KEGG pathway enrichment analysis of genes with significantly altered splicing events. The top 5 enriched terms from each group are shown (adjusted P < 0.05, Benjamini–Hochberg correction). Numbers indicate the count of genes from the input list associated with each enriched term. Representative gene symbols contributing to the enrichment of each category are listed below the term name.

Because Ilf2 has been implicated in RNA processing^24^, we next examined alternative splicing. Bulk RNA-seq revealed widespread splicing reprogramming, predominantly skipped-exon events (**Figure 3h and i, Supplementary Table S7**), with clear exon-usage shifts at functionally relevant loci, exemplified by Cttn^25^ (ENaC regulation in collecting-duct epithelia) and Fyn^26–30^ (renal fibrosis, diabetic kidney injury, and hyperosmotic stress **Figure 3j)**. Differentially spliced genes were enriched in pathways associated with cellular adaptive responses, including cytoskeleton^31^ and junctional architecture^32^ (e.g., actin-filament organization, growth cone, adherens / tight junction), as well as ATP-dependent chromatin remodeling^33^ **(Figure 3k, Supplementary Table S7)**.

Together, these orthogonal data indicate that Ilf2 coordinates transcriptional and post-transcriptional programs that underpin intrinsic epithelial resilience.

### Ilf2 sustains proliferation and stress resilience of collecting duct cells

Guided by the transcriptomic predictions, we tested functional consequences of Ilf2 loss in IMCD3 cells. Ilf2 knockdown led to a modest but significant reduction in proliferation, reflected by fewer Ki67-positive cells **(Figure 4a–b)**. Moreover, Ilf2-deficient cells exhibited abnormal nuclear morphology, with increased frequencies of multinucleated or irregularly shaped nuclei **(Figure 4c–d)**, indicating defects in mitotic division and nuclear integrity. Under hyperosmotic challenge (600 mOsm), Ilf2 deficient cultures showed reduced viability and higher death ratios relative to controls **(Figure 4e-g)**, demonstrating impaired tolerance to osmotic stress. Collectively, these findings establish that Ilf2 supports proliferation, nuclear stability, and osmotic-stress tolerance in IMCD3 cells.

**Figure 4.**
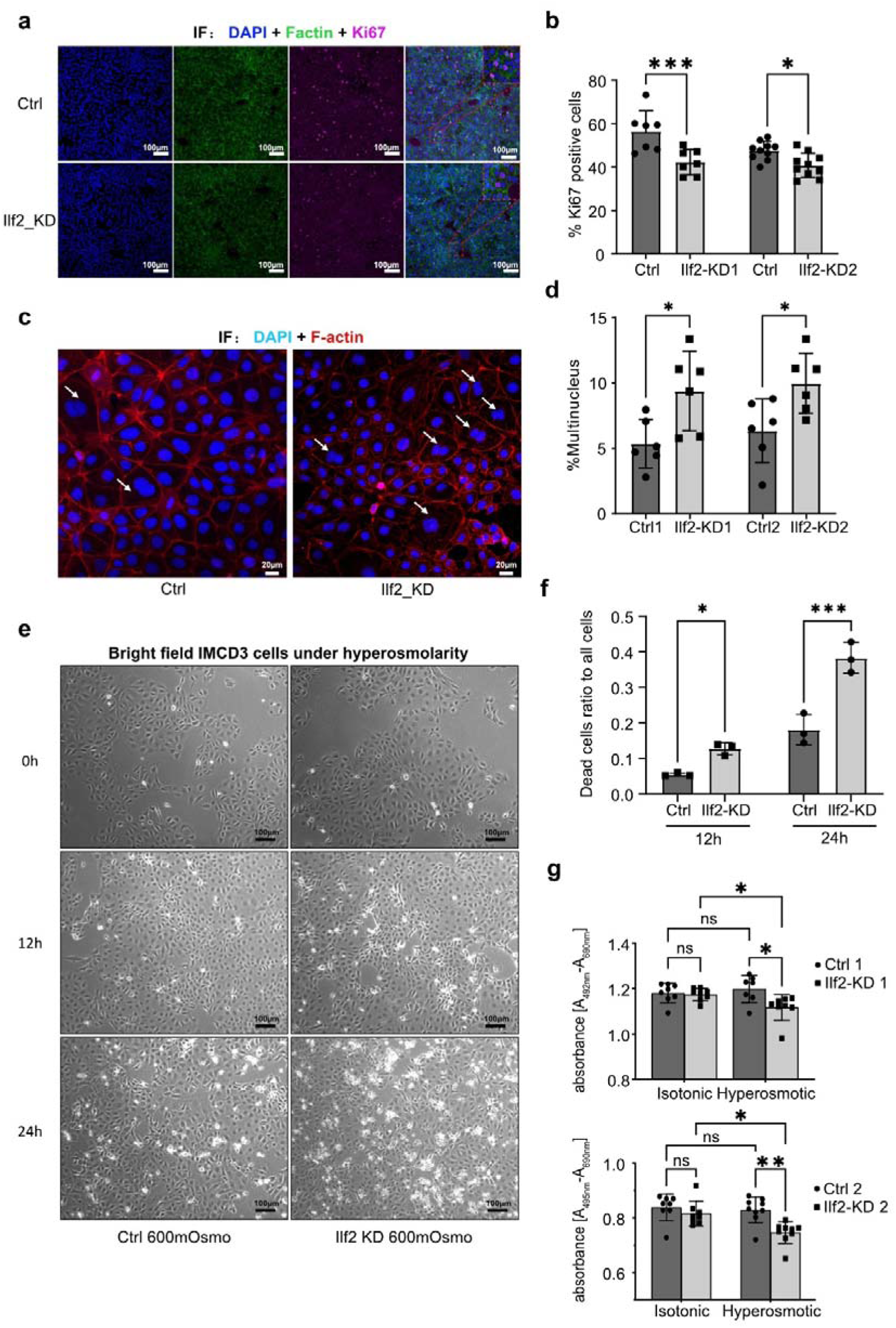
Ilf2 biological roles in IMCD3 cells. **(a):** Ki67 staining in cells transduced with two different Ilf2-targeting gRNAs (Ilf2 knockdown, KD1–2) and two non-targeting control gRNAs (Ctrl1–2). DAPI (blue) was used as a nuclear counterstain, Ki67 was labeled in pink, and F-actin (green) was used to delineate cell boundaries. **(b):** Quantification of Ki67 positive cells ratio in one culture well, each dot represents an independent well (gRNA pair 1 n=7, gRNA pair 2 n=10). Statistical significance between control and knockdown groups was assessed using two-way ANOVA followed by Bonferroni-corrected pairwise comparisons. **(c):** DAPI staining of the Ilf2 knockdown cells and control cells, F-actin was stained for the cell border detection. Multi-nucleus cells were pointed out with arrows. **(d):** Quantification of the multi-nucleated cells, each dot represents an independent culture well. The analysis included two different gRNA pairs (gRNA pair 1 n=4, gRNA pair 2 n=4). Statistical significance between control and knockdown groups was assessed using two-way ANOVA followed by Bonferroni-corrected pairwise comparisons. **(e):** Representative phase-contrast images of Ilf2 knockdown and control IMCD3 cells after 12-hour exposure to 600 mOsm hyperosmotic medium. Dying cells exhibit characteristic morphological changes, including a rounded shape, increased cytoplasmic refringence (bright border), and detachment from the culture surface^34^. **(f):** Quantification of dead cells relative to living cells was performed in ImageJ. Each dot represents an independent well (n=3 for each time point per condition). Statistical significance between control and knockdown groups was assessed using two-way ANOVA followed by Bonferroni-corrected pairwise comparisons. **(g):** Quantification of cell viability under isotonic or hyperosmotic conditions by measuring the absorbance with XTT assay. Each dot represents an independent well (2 pairs of gRNA were quantified separately, gRNA pair 1 n=8, gRNA pair 2 n=8). Tukey’s multiple comparisons test was used to access the statistical significance. * adjusted *P* < 0.05; ** adjusted *P* < 0.01; *** adjusted *P* < 0.001.

### Ilf2 is enriched in collecting duct and activated after ischemia–reperfusion injury in mouse

To connect these phenotypes with *in vivo* expression, we analyzed Ilf2 distribution in mouse kidney. snRNA-seq and bulk RNA-seq revealed preferential Ilf2 expression in collecting duct cells of the inner medulla **(Figure 5a-b)** consistent with a re-analysis of bulk RNA-seq data of micro-dissected mouse kidney^35^ **(Figure 5c).** *In situ* hybridization and immunofluorescence confirmed strong Ilf2 signals in Aqp4- and Aqp2-positive cells in the medulla, whereas weaker Ilf2 signals were detected in other medullary and cortical cells (**Figure 5d)**. These data validated that Ilf2 is selectively enriched in the renal medullar, particularly in medullary collecting duct cells, but also expressed in several other kidney cell types.

**Figure 5.**
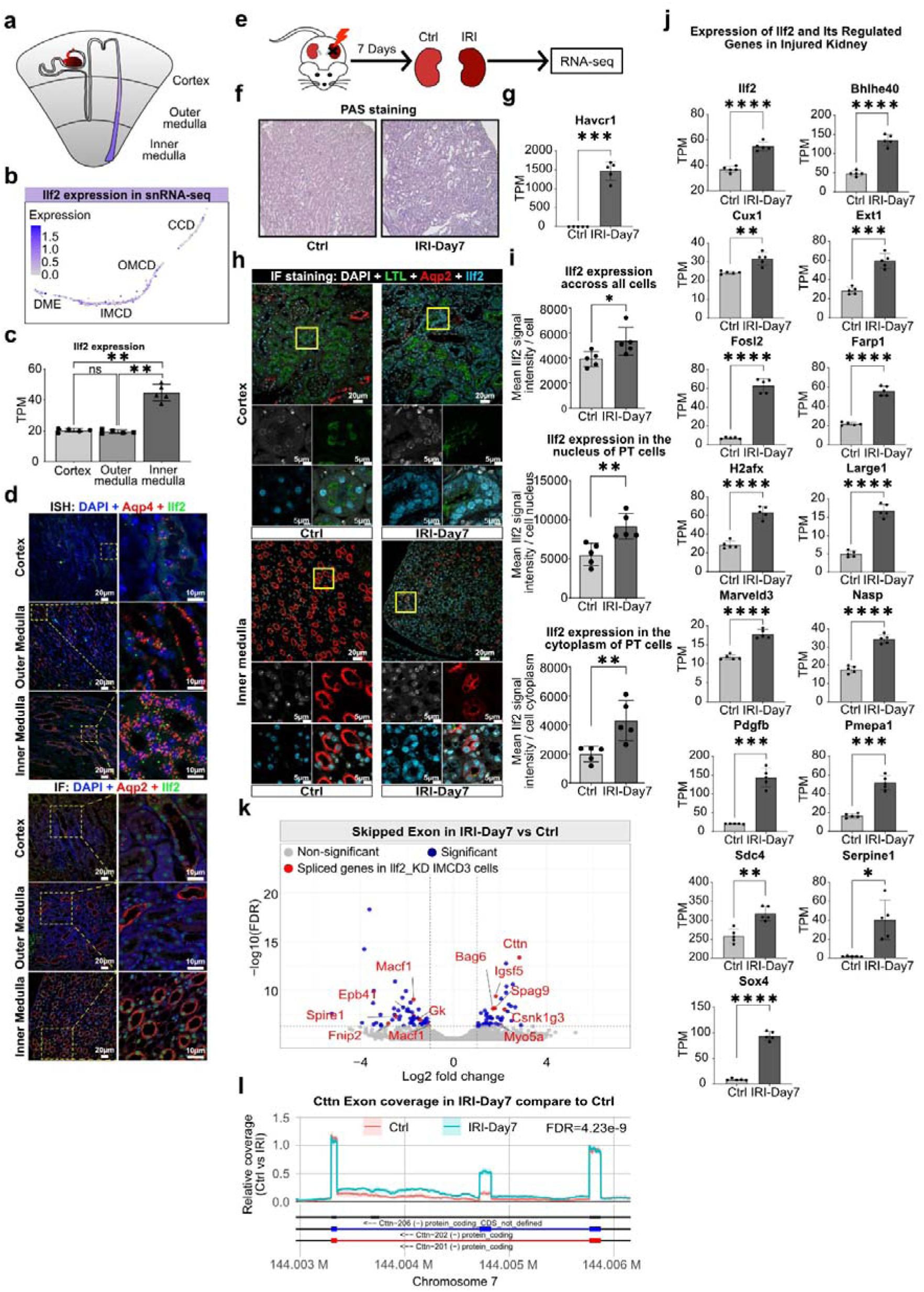
Ilf2 expression *in vivo* and it’s coordinated gene expression in IRI. **(a):** Schematic illustration of nephron tubules and the RNA expression level of the Ilf2 as shown in purple. **(b):** sn-seq UMAP plot of sub-clustered collecting duct cells, featuring the expression of the Ilf2 gene, the expression level was log-normalized with the Seurat data processing package. **(c):** Bulk RNA-seq data from cortex, outer medulla, and inner medulla regions of the mouse kidney were reanalyzed to assess Ilf2 expression levels. The data were obtained from a previously published study by our group^35^. The expression was normalized to transcripts per million (TPM). Statistical significance was assessed using Welch’s ANOVA with Brown–Forsythe correction, appropriate for datasets with unequal variances across groups. NS: not significant (adjusted *P* > 0.05), ** adjusted *P* < 0.01. **(d; upper panel)** In situ hybridization (ISH) was performed on 2 WT mouse kidney slices (n=2) to visualize Ilf2 mRNA expression. Samples were stained with DAPI and labeled for Aqp4 (A marker of collecting duct principal cells) and Ilf2. Representative images from the cortex, outer medulla, and inner medulla regions are displayed to illustrate the distribution of Ilf2 mRNA levels within the kidney collecting duct. Zoomed-in images of selected regions were supplied on the right side. **(d; lower panel):** Immunofluorescence (IF) staining was performed on 2 WT mouse kidney slices (n=2) to visualize Ilf2 protein expression. Samples were stained with DAPI, mouse anti-Aqp2 (A marker of collecting duct principal cells), and mouse anti-Ilf2. Representative images from the cortex, outer medulla, and inner medulla regions are displayed to illustrate the distribution of Ilf2 protein levels within the kidney collecting duct. Zoomed-in images of selected regions were supplied on the right side. **(e):** Schematic representation of the experimental workflow, unilateral ischemia reperfusion injury (uIRI) was utilized, contralateral (Ctrl) and injured (IRI) kidney samples were sequenced (bulk RNAseq) 7 days post-injury. **(f):** Representative kidney sections from Ctrl and IRI mice at day 7 post-injury were stained with PAS to assess tubular morphology and basement membrane integrity. **(g):** Kidney injury marker gene (Havcr1) expression at Day7. Each dot represents an individual mouse, n=6 per group. Statistical significance was assessed using a t-test with Welch’s correction between Ctrl and IRI samples. **(h):** Immunofluorescence staining was performed on kidney sections from 5 contralateral control (Ctrl) and 5 ischemia-reperfusion injury (IRI) mice (n = 5 per group) to assess Ilf2 protein expression. Sections were stained with DAPI (nuclei), mouse anti-Aqp2 (collecting duct marker), mouse anti-Ilf2, and Lotus Tetragonolobus Lectin (LTL, proximal tubule brush border marker). Representative images from the cortex and medulla are shown. Under IRI conditions, increased Ilf2 expression and subcellular redistribution from the nucleus to the cytoplasm were observed. Insets highlight Ilf2 translocation in magnified views. **(i):** Quantification of Ilf2 expression across all cell types and within the nucleus and cytoplasm of proximal tubule (PT) cells. The signal intensity of individual cells was measured using QuPath software. For each sample, the mean signal intensity was calculated across all detected cells or specifically within PT cells. The cytoplasmic region was defined as a 5 µm-wide area surrounding each identified nucleus. Each dot represents one independent kidney sample. Statistical significance was assessed by Mann-Whitney U test. **(j):** Genes downregulated upon Ilf2 knockdown exhibit upregulation in the injured kidney, including Ilf2. Each dot represents an individual mouse, n=6 per group. Statistical significance was assessed using a t-test with Welch’s correction between Ctrl and IRI samples. **(k):** Volcano plots showing differentially spliced events in skipped exons (SE) category in IRI compared to control kidneys. The x-axis indicates log_₂_ fold change in splicing event inclusion; the y-axis indicates –log_₁₀_(FDR). Red dots and gene name (red) highlight splicing events that overlap with those observed in Ilf2-knockdown IMCD3 cells (compare to Fig. 6). Multiple testing correction was performed using the Benjamini–Hochberg correction to determine statistical significance. In the plot, genes with an adjusted p-value < 0.05 and a log_₂_ fold change > 1 were considered significant. **(l):** Representative alternative spliced gene Cttn inclusion in IRI samples. Coverage plots display normalized read densities across the Cttn (chr7:144.003–144.006 Mb) gene loci. Each curve represents the mean normalized coverage across replicates for each condition: Ctrl (red) and IRI (blue). Shaded areas indicate the standard error. Transcript isoforms are shown below with exon–intron structures. * adjusted *P* < 0.05; ** adjusted *P* < 0.01; *** adjusted *P* < 0.001, **** adjusted *P* < 0.0001.

Given that Ilf2 loss renders IMCD3 cells maladaptive under osmotic stress **(Figure 4e)**, and ischemia–reperfusion injury (IRI) is a canonical renal stress paradigm^36^, we hypothesized that kidney injury would induce Ilf2 and engage Ilf2-dependent programs *in vivo*. Although ischemia–reperfusion injury (IRI) predominantly affects proximal tubules^4^, collecting duct cells are likewise exposed to the injury milieu and contribute to epithelial recovery^37^. We therefore employed a unilateral IRI model **(Figure 5e-f)**. In this context, Ilf2 expression was markedly upregulated by day 7 post-injury, as confirmed by RNA-seq and immunostaining **(Figure 5h–j)**. Importantly, this upregulation was not confined to collecting ducts, but affected additional cortical and medullary cell types. Concordantly, numerous Ilf2-regulated genes that were suppressed in Ilf2-KD IMCD3 cells—including Bhlhe40, Fosl2, H2afx, and Serpine1—were upregulated during kidney repair **(Figure 5j)**. These reciprocal patterns support a model in which Ilf2 orchestrates transcriptional programs that promote stress adaptation and potentially participate in tissue recovery.

We also observed parallels at the level of RNA splicing. Alternative splicing events perturbed by Ilf2 knockdown *in vitro* were inversely regulated in IRI kidneys. For example, Cttn displayed increased exon inclusion in post-ischemic kidneys, whereas Ilf2 depletion induced exon skipping **(Figure 5k–l).** These findings suggest that Ilf2 activity *in vivo* directs splicing programs that favor cytoskeletal remodeling and stress resistance.

Notably, Ilf2 changed its subcellular distribution after IRI. While largely nuclear in uninjured kidneys, Ilf2 accumulated in the cytoplasm of post-injury tubular cells **(Figure 5h–i)**. Given that NF45/Ilf2 can shuttle to the cytoplasm and support selective translation under stress^38,39^, this cytoplasmic shift might enable synthesis of pro-survival proteins when global translation is impaired. Together, these results demonstrate that Ilf2 is activated during kidney injury and orchestrates transcriptional and splicing programs that enhance epithelial survival and repair.

### ILF2 is increased in human kidneys with advanced CKD

To determine whether the ILF2 upregulation observed in the mouse kidney injury model is conserved in human kidney disease, we performed immunofluorescence staining for ILF2 in renal biopsies from patients with mild or advanced forms of chronic kidney disease (CKD) that underwent kidney biopsies **(Table 1)**. Samples represented different etiologies of kidney disease, including different glomerular diseases and post-transplant injuries. Patients classified as advanced CKD displayed extensive tubulointerstitial fibrosis and tubular atrophy (IFTA > 30%) and displayed reduced glomerular filtration rate (eGFR ≤ 30 ml/min/1.73 m2), whereas those classified as mild CKD displayed substantially less IFTA (≤ 5%) and exhibited preserved kidney function (eGFR >90 ml/min/1.73 m2). In patients with mild CKD, ILF2 signals were predominantly nuclear in tubular epithelial cells, including proximal tubules (PT) and collecting ducts (CD) **(Figure 6a**). In contrast, in patients with advanced CKD, quantitative analysis of ILF2 signal intensity revealed significantly enhanced nuclear ILF2 signals both in PT and CD cells when compared to patients with mild injury **(Figure 6a and b)**. Consistently, ILF2 intensity in PTs correlated positively with serum creatinine levels (r = 0.68, P = 0.025; **Figure 6c**) and negatively with eGFR (r = −0.72, P = 0.012; **Figure 6d**). In addition, ILF2 intensity in PT tended to correlate positively with the degree of interstitial fibrosis (r = 0.59, P = 0.059; **Figure 6e**). Interestingly, patients with advanced CKD displayed cytoplasmic ILF2 staining in PT and CD cells **(Figure 6a)**, similar to what was observed in injured mouse kidneys **(Figure 5h)**. These findings indicate that ILF2 expression in renal tubules is consistently enhanced in advanced CKD, consistent with a stress-responsive upregulation.

**Figure6.**
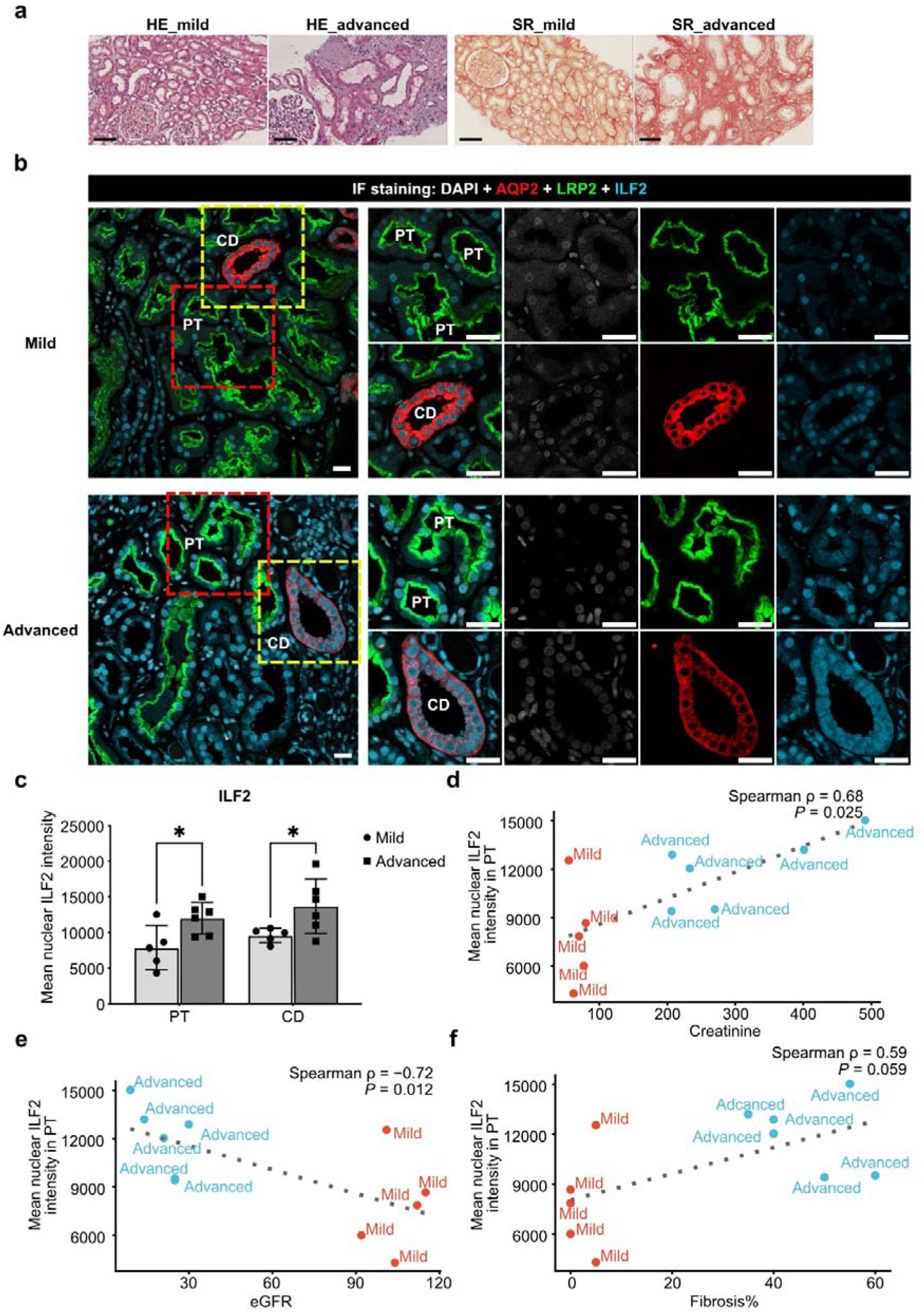
ILF2 is upregulated in human kidneys with advanced CKD. **(a)** Representative histological images of human kidney sections from patients with mild and advanced chronic kidney disease (CKD). Hematoxylin and eosin (HE) staining shows tubular atrophy and glomerulosclerosis in advanced CKD, while Sirius Red (SR) staining highlights increased collagen deposition and interstitial fibrosis. Scale bars, 100 µm. **(b)** Immunofluorescence images of human kidney biopsies (mild vs. advanced CKD). Representative fields stained for DAPI (nuclei, white), LRP2 (proximal tubule [PT] marker, green), AQP2 (collecting duct [CD] principal-cell marker, red), and ILF2 (cyan). Top row: Mild CKD; bottom row: Advanced CKD. The first column shows the merged view. Red dashed boxes highlight PT-rich areas; yellow dashed boxes highlight CD-rich areas. Columns to the right show zoom-ins of the boxed regions followed by single-channel images for LRP2, AQP2, and ILF2, respectively. PT and CD labels indicate tubular segment identity based on marker expression. Scale bars in zoomed in images are 20µm. **(c)** Mean nuclear ILF2 intensity in PT and CD per patient, comparing mild vs advanced CKD. Bars represent mean ± SD.; each data point is one patient. Asterisks indicate statistically significant differences between mild and advanced CKD (Turkey’s multiple comparison test with correction). Quantification was performed on nuclei detected within LRP2^⁺^ (PT) or AQP2^⁺^ (CD) regions using QuPath software. **(d-f)** Clinical correlations of PT nuclear ILF2 intensity. Scatter plots showing per-patient PT nuclear ILF2 intensity vs clinical/pathology readouts. Points are annotated by group (Mild vs Advanced CKD). Statistics shown in each panel are Spearman correlation coefficients (ρ) with two-sided *P* values. **(d)** Positive correlation with serum creatinine. **(e)** Negative correlation with eGFR. **(f)** Positive correlation with IFTA (% value). * adjusted *P* < 0.05.

**Table 1:**
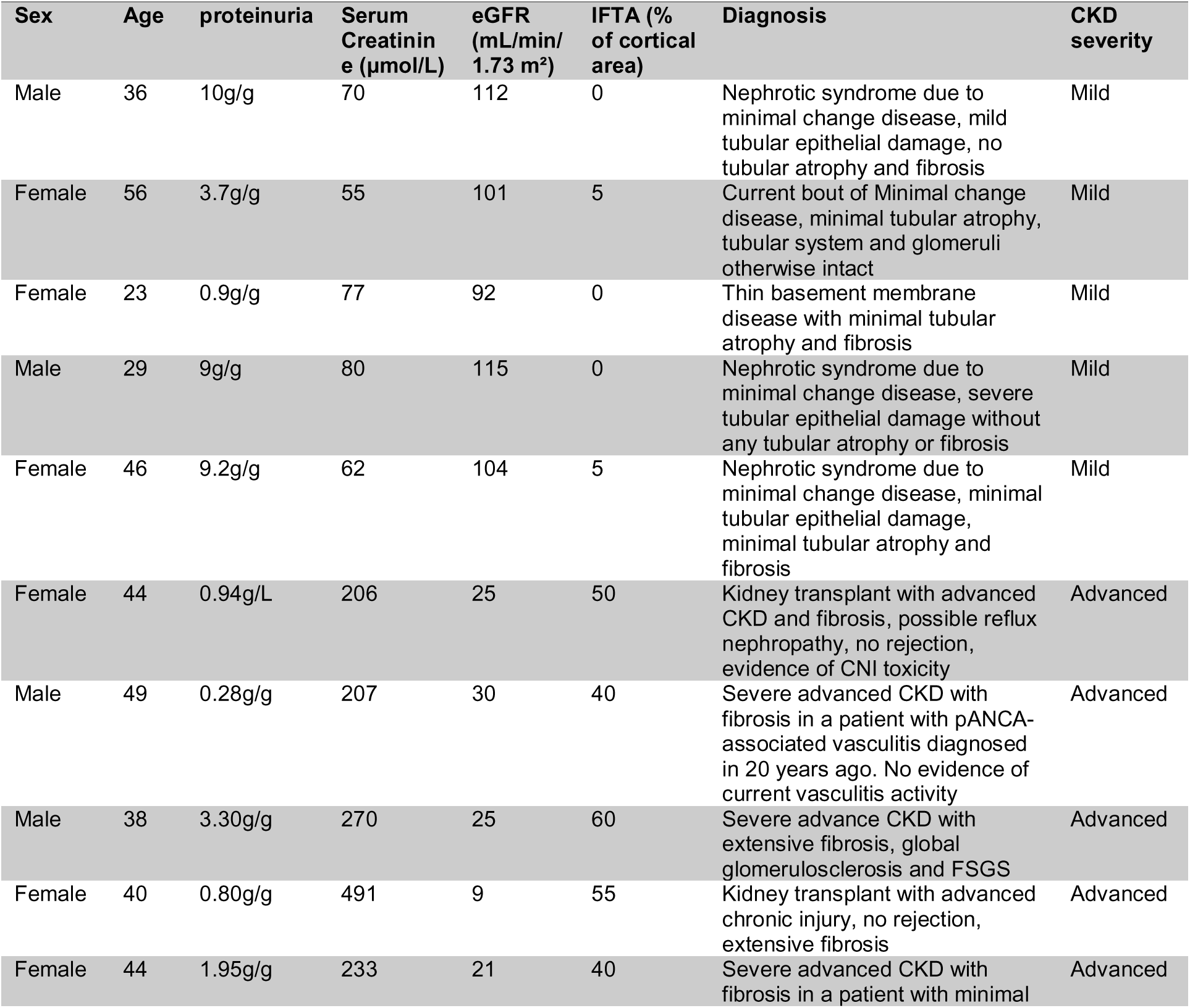
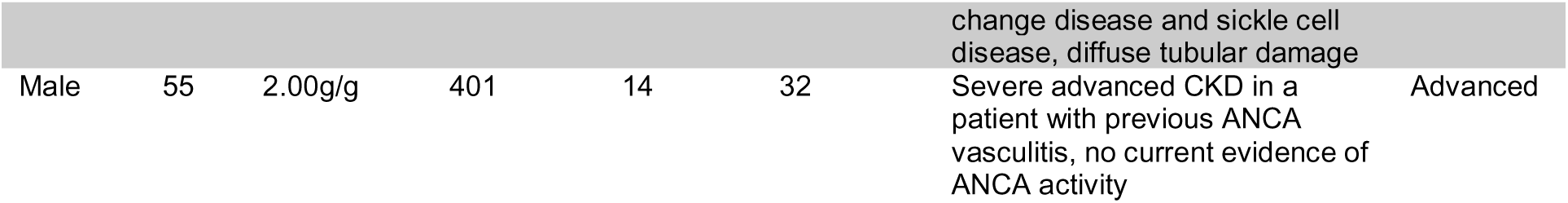
Clinical and pathological features of patient cohort.

## Discussion

By integrating single-cell functional genomics and *in vivo* validation, we have identified Ilf2 (NF45) as a transcriptional regulator that promotes epithelial stress resilience in the kidney. Unlike previously studied TFs such as Grhl2, Ilf2 was not previously linked to kidney tubular functions or injury. Our study expands Ilf2’s functional repertoire (known mostly from cancer biology and immunology) to include renal tubular integrity, cytoskeletal organization, and stress-adaptive RNA processing.

A major challenge in molecular biology is identifying and characterizing “hidden” regulators that classical approaches might overlook. We addressed this by combining bioinformatics-driven candidate selection with high-throughput single-cell CRISPR screening, allowing us to functionally test many TFs in parallel. This strategy efficiently pinpointed Ilf2 as a necessary regulator among numerous candidates. The Perturb-seq approach proved powerful – it not only confirmed known regulators (e.g., Grhl2, Klf5 and Junb) but also unveiled Ilf2’s impact. This underscores the value of single-cell CRISPR screens in uncovering complex regulatory networks in specialized cell types.

Ilf2/NF45 is a multifunctional RNA-binding protein. Past studies in other systems show Ilf2 (NF45) partners with Ilf3 (NF90/NF110) to regulate cell cycle progression, mitosis, ribosome biogenesis, and RNA splicing^24,40–46^. We extend these findings by showing that Ilf2 integrates those activities into a stress resilience context in kidney epithelium. For medullary cells constantly under osmotic and hypoxic stress, Ilf2 appears to calibrate gene expression to maintain structural components (cytoskeleton, junctions) and ensure proper mRNA maturation for stress-response genes.

One particularly novel aspect is the interplay between Ilf2 and alternative splicing of stress-related genes (like Fyn and Cttn). The prevalence of skipped exon events upon Ilf2 loss indicates Ilf2 helps include certain exons – potentially those important for full protein functionality. For example, Fyn kinase’s role in fibrosis^26,27^ and osmoadaptation^47^ might depend on specific isoforms that Ilf2 helps generate. Similarly, the Cttn variant favored by Ilf2 could strengthen actin networks^48^, aiding cell survival in high osmolarity^25^. NF45/NF90/NF110 complexes have been shown to associate with splicing machinery^24^; our findings suggest that in renal cells under stress, Ilf2 ensures correct splicing of key structural and signaling genes, thereby reinforcing cell stability and signaling capacity.

The *in vivo* data from the IRI model and human mild and advanced CKD samples provide a physiological context: during injury and recovery, Ilf2 is induced along with many of its target genes. The upregulation of Ilf2 and Ilf2-dependent transcripts (e.g., H2afx for DNA damage response^49,50^, Serpine1 for tissue remodeling^51,52^, Bhlhe40 for hypoxia adaptation^53^) suggests a coordinated stress response program. This coordination implies Ilf2 could be part of an intrinsic protective response – as the kidney enters a repair phase, Ilf2 levels rise to drive expression of genes that facilitate recovery and protect against further stress.

Intriguingly, Ilf2 translocates to the cytoplasm in injured cells, possibly to enable selective translation. Cytoplasmic Ilf2/NF45 has been reported to enhance internal ribosome entry site (IRES)-mediated translation of anti-apoptotic proteins during ER stress^38^. We speculate a similar mechanism in post-ischemic kidneys: Ilf2 might help synthesize survival proteins at a time when global protein synthesis is inhibited. This would allow stressed tubular cells to produce critical proteins (like chaperones or cell-cycle regulators) despite general translational shutdown. Such a role would elegantly complement Ilf2’s nuclear function – bridging transcriptional control and translational control in an integrated stress response. While further experiments are needed to confirm this cytoplasmic function in kidneys, it represents a plausible link between our *in vitro* and *in vivo* observations.

In summary, our data demonstrate that Ilf2/NF45 is a central regulator that ties together transcriptional and splicing programs to reinforce epithelial resilience under stress. This is particularly relevant to the challenging environment of kidney epithelia under conditions of hyperosmolality, hypoxia or oxidative stress. The induction of Ilf2 in kidney injury and its dynamic localization suggest it actively participates in injury responses, repair processes, and resilience to future insults. Thus, Ilf2 might serve as both a biomarker of a cell’s stress-adaptive state and a potential therapeutic target. Augmenting Ilf2 function (or downstream effectors) could hypothetically boost the stress tolerance of tubular cells, thereby reducing injury in AKI or enhancing recovery. Conversely, inhibiting Ilf2 might be detrimental in AKI but could potentially be explored in contexts where limiting cell proliferation is desired (e. g. in renal neoplasia). Our findings open new avenues to explore stress resilience pathways in epithelia, moving closer to interventions that protect the kidney from acute and chronic insults.

## Data Sharing Statement

Single-cell, single-nuc and bulk RNA sequencing data have been deposited to Gene Expression Omnibus (GEO), accession numbers: GSE299239 (reviewer token onwzwycqtvmntgl), GSE299244 (reviewer token gxibwcquhngpnml), GSE299248 (reviewer token qtklsmwqffwppsz), GSE299251 (reviewer token GSE299251), and GSE299249 (reviewer token edutiqmqxhqxnqp). The primary scripts used for data analysis in this study are publicly available on GitHub at https://github.com/shuangcao-mhh/Perturb-seq-screening-in-IMCD3-cells. Additional research data underlying this work is deposited at the MHH research data server https://doi.org/10.26068/mhhrpm/20250530-000 and can be made available upon request.

## Acknowledgments

This project was supported by grants awarded to K.M.S.O: German Research Foundation (DFG) Research Training Group GRK 2318 (B4) – Project ID 318905415. We thank Tatjana Luganskaja, Martina Flechsig, and Gabriele N’Diaye for excellent technical support.

## Author contributions

SC and KMSO designed the study; SC, KILC, JL and FJB conducted experiments; SC, CH analyzed data; JS and JHB provided and prepared human samples; SC made the figures; SC and KMSO drafted and edited the manuscript; all authors edited the manuscript and approved the final version of the manuscript.

