## Supplementary Data for "Interleukin Enhancer Binding Factor 2 (Ilf2) and Kidney Epithelial Stress Resilience"

**SUPPLEMENTAL CONTENT:**

**Supplementary Figure S1.** Generation of dCas9-KRAB IMCD3 cell lines.

**Supplementary Figure S2.** Optimization of four CRISPRi single-cell compatible vectors.

**Supplementary Figure S3.** Quality control metrics for single-cell RNA-seq data across targeted gRNAs.

**Supplementary Figure S4.** Correlation between gRNA library abundance and single-cell coverage across target genes.

**Supplementary Tables (Supplementary tables S1-S7 are provided as a single Excel file).**

**Supplementary Table S1.** Candidate transcription factors selected for CRISPRi screening.

**Supplementary Table S2.** Cell type-based regulon activity scores and regulon-specific scores (Excel file with two sheets: “activity” and “specificity”).

**Supplementary Table S3.** Normalized gene expression from bulk RNA-seq of WT IMCD3 cells.

**Supplementary Table S4.** Pseudo-bulk analysis of differentially expressed genes following knockdown of Grhl2, Klf5, Junb, and Ilf2.

**Supplementary Table S5.** Functional enrichment analyses following knockdown of Klf5, Junb, and Ilf2.

**Supplementary Table S6.** TPM-normalized gene expression (for visualization) and DESeq2-based differential expression analysis of Ilf2 knockdown IMCD3 cells, with ComBat-seq batch correction applied to raw counts.

**Supplementary Table S7.** Differentially spliced events following Ilf2 knockdown and enrichment analysis of associated genes.

**Supplementary Figure S1**

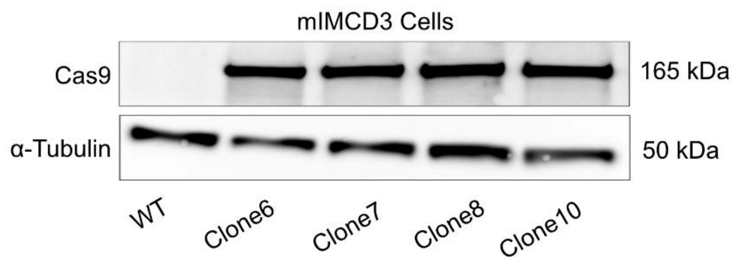


**Supplementary Figure S1: Generation of dCas9-KRAB IMCD3 cell lines.** Western blot analysis showing stable expression of dCas9-KRAB in four selected clones (clones 6, 7, 8, and 10) compared to wild-type (WT) IMCD3 cells. Cas9 expression (~165 kDa) was detected using a specific antibody, with α-Tubulin serving as a loading control.

### Supplementary Figure S2

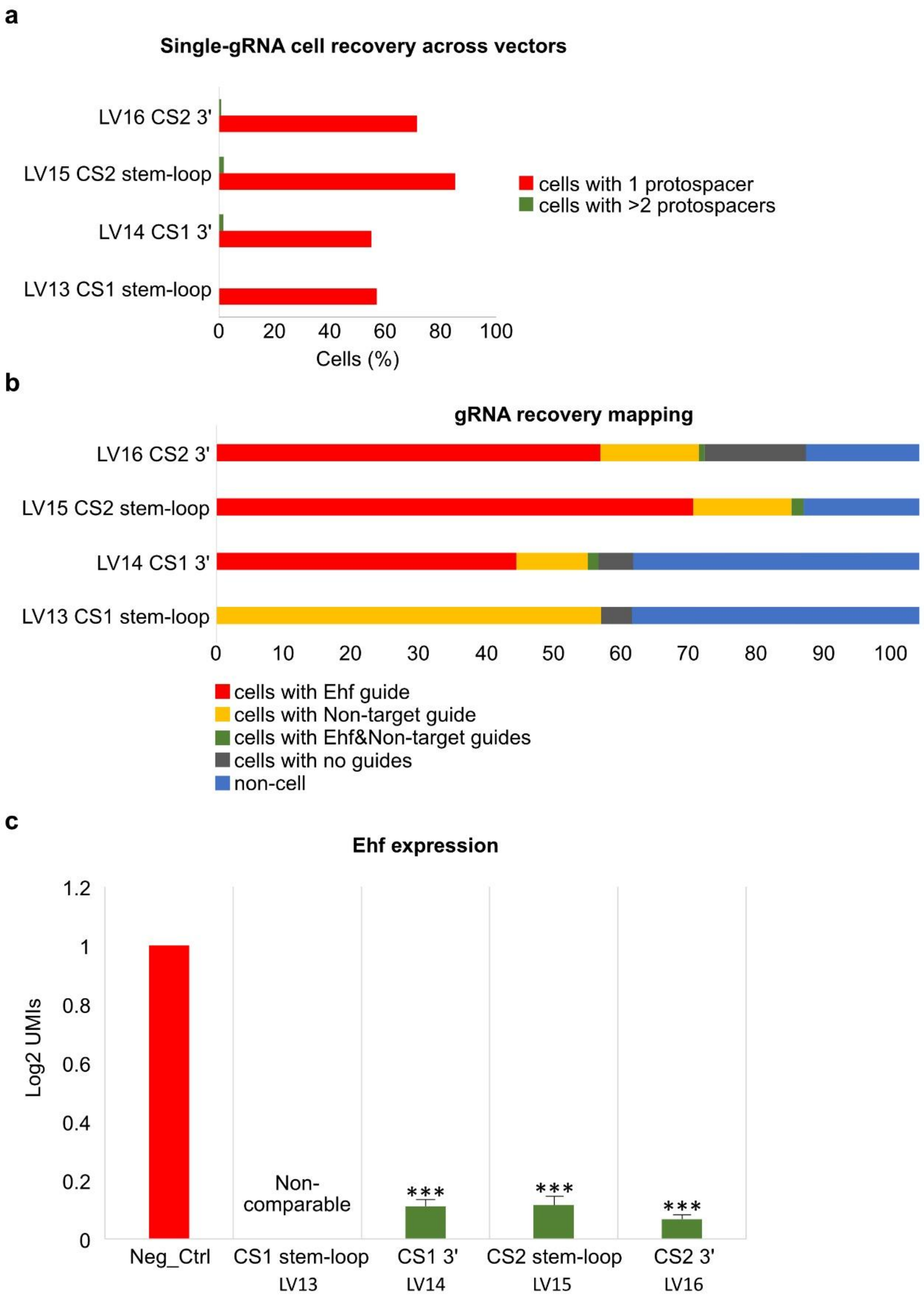


**Supplementary Figure S2: Optimization of four CRISPRi single-cell compatible vectors.** Four gRNA delivery vectors were tested in IMCD3 cells, each incorporating a capture sequence (CS1 or CS2) either within the gRNA stem-loop or at the 3’ end: LV13 (CS1 stem-loop), LV14 (CS1 3’), LV15 (CS2

stem-loop), and LV16 (CS2 3’). Each vector encoded one gRNA targeting the mouse gene Ehf and one non-targeting control gRNA. **a:** Quantification of single-gRNA cell recovery across vectors. Red bars represent the percentage of cells with a single gRNA (1 protospacer), and green bars represent cells with more than one gRNA (>2 protospacers). **b:** gRNA recovery mapping. Stacked bars show the proportion of cells carrying Ehf guides (red), non-targeting guides (yellow), both (gray), no guides (blue), or classified as non-cell (light blue). **c:** Ehf expression following knockdown using each vector. Expression is shown as $\log_2$ UMIs. A negative control (Neg_Ctrl) was included. Multiple testing correction was performed using the Benjamini–Hochberg (BH) method. *** indicates adjusted $P < 0.001$.

### Supplementary Figure S3

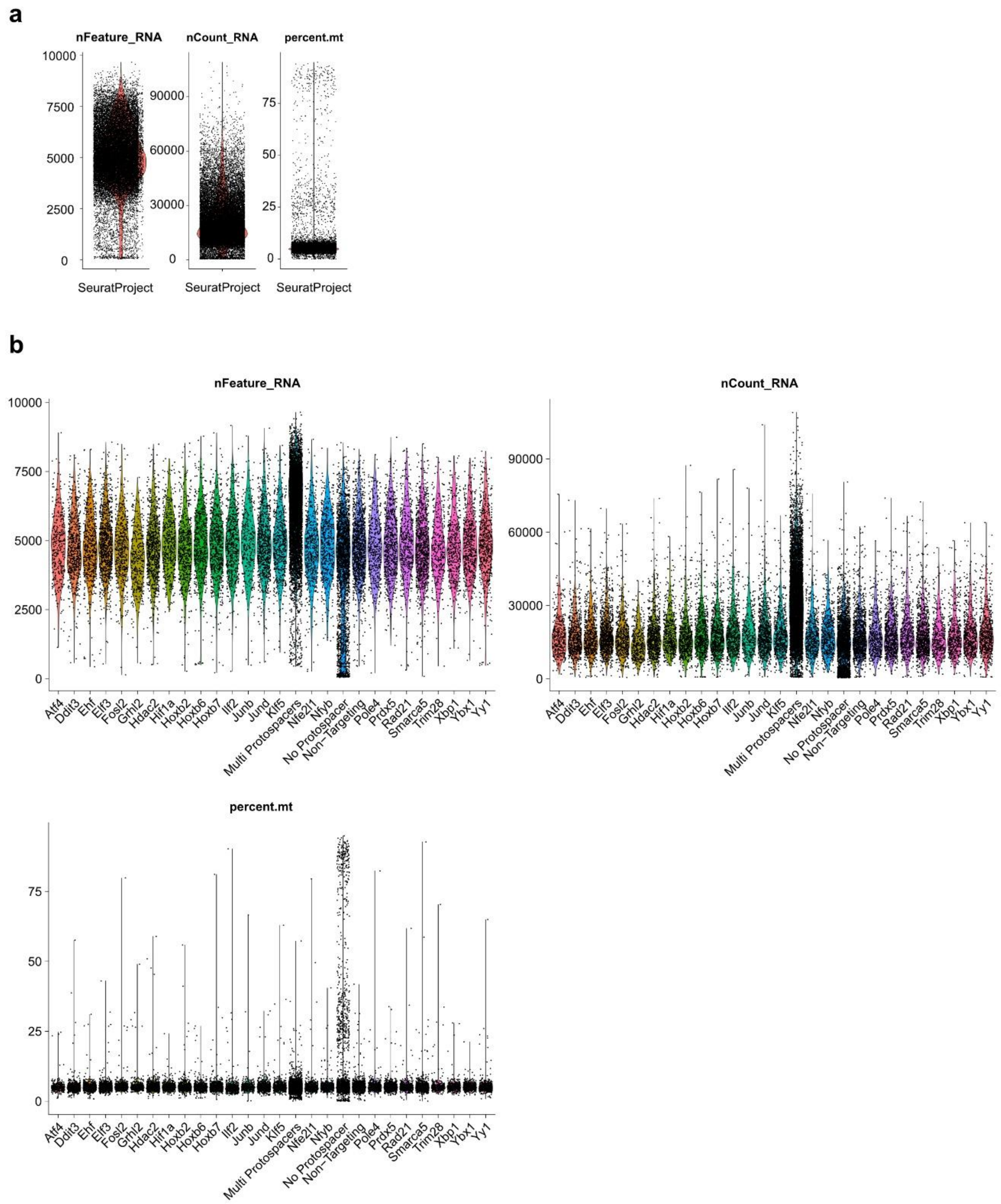


**Supplementary Figure S2: Quality control metrics for single-cell RNA-seq data across targeted gRNAs. a:** Violin plots showing the distribution of the number of detected genes per cell (nFeature_RNA), total RNA counts per cell (nCount_RNA), and the percentage of mitochondrial transcripts (percent.mt) across all cells before filtering. **b:** Violin plots showing the distribution of detected genes per cell (nFeature_RNA), total RNA counts per cell (nCount_RNA), and the percentage of mitochondrial transcripts (percent.mt), grouped by sample identity based on gRNA assignment. The No

Protospacer group, which contains cells lacking assigned gRNAs, displays elevated mitochondrial content, likely reflecting low cell quality. As these cells lack target gRNA information, they were excluded from downstream analyses.

**Supplementary Figure S4**

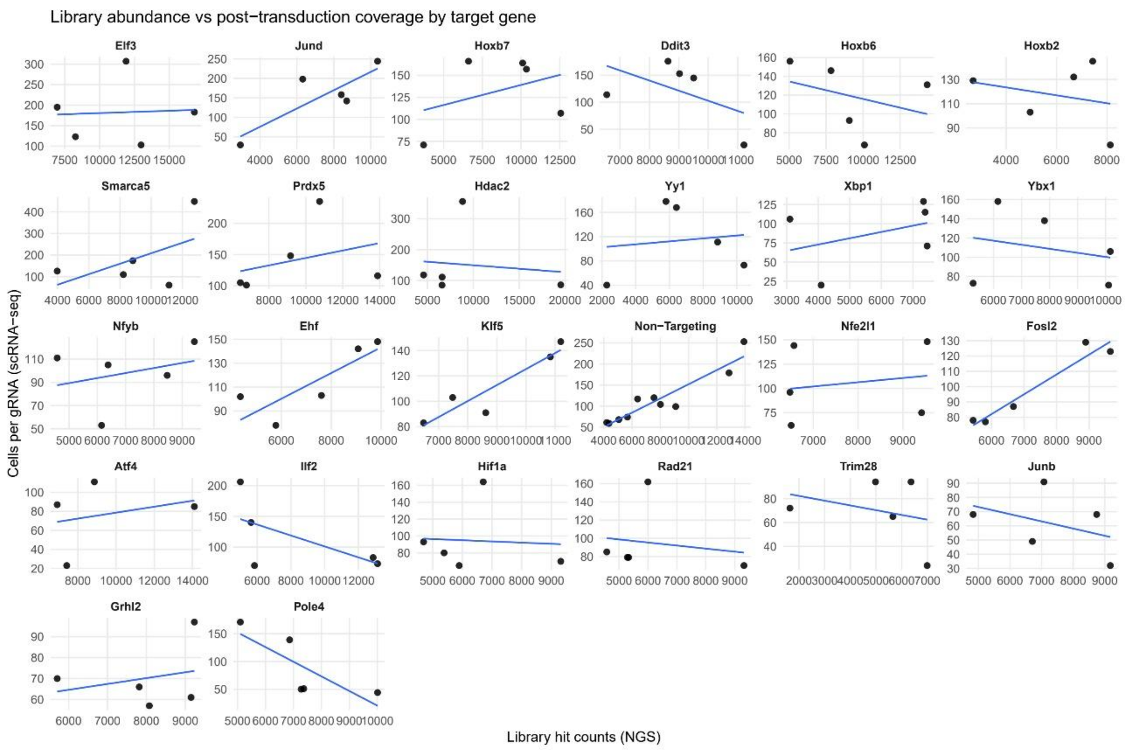


**Supplementary Figure S4: Correlation between gRNA library abundance and single-cell coverage across target genes.**

Scatter plots show the relationship between the lentiviral library abundance of each gRNA (x-axis, quantified by next-generation sequencing, NGS) and the number of cells carrying that gRNA detected by single-cell RNA-seq (y-axis). Each panel represents a different target gene included in the CRISPRi library. Blue lines indicate linear regression fits. Overall, gRNA representation in scRNA-seq was consistent with the initial library distribution, confirming efficient gRNA capture and unbiased transduction.
